# Testing monotonic trends among multiple group means against a composite null

**DOI:** 10.1101/2020.05.28.120683

**Authors:** Chenxiao Hu, Thomas Sharpton, Duo Jiang

**Affiliations:** Department of Statistics, Oregon State University; Department of Microbiology, Oregon State University

## Abstract

In biomedical applications, it is often of interest to test the alternative hypothesis that the means of three or more groups follow a strictly monotonic trend such as *µ*_1_ *> µ*_2_ *> µ*_3_ against the null hypothesis that the group means are either equal or unequal but are not monotonic. This is useful, for example, for detecting biomarkers whose level in healthy, low-risk cancer and aggressive cancer subjects increases or decreases throughout the three groups. Various trend tests are available for testing monotonic alternatives. However, existing methods are designed for a highly restrictive null hypothesis where all group means are equal, which represents a special case of the null space in our problem. We demonstrate that these methods fail to control type I error when the group means may be unequal under the null. To test this broader null hypothesis, develop a greedy testing method which has an intuitive interpretation related to two-sample t tests. We show both theoretically and through simulations that the proposed method effectively controls type 1 error throughout the entire null space and achieves higher power than a naive implementation of multiple t-tests. We illustrate the greedy trend test method in real data to study microbial associations with parasite-related pathology in zebrafish.

## 1 Introduction

Statistical testing of equality of population means among three or more groups has been studied for decades. Familiar techniques including analysis of variance and non-parametric methods such as the Kruskal-Wallis test are widely used. In many applications [1, 2], it is of interest to focus on a more specific alternative hypothesis where the group means follow a monotonic order. Various approaches are available to test for monotonic trends among the group means. For binary data, the Cochran-Armitage test[3, 4] is a modification of the Pearson Chi-square test to detect association among multiple groups. For continuous data, under some assumptions, linear regression may be performed by including the group label *i* as an explanatory variable in order to detect a linear trend between the group means. There are also nonparametric methods such as rank-based tests for the monotonic ordering problem. Examples include the Jonckheere-Terpstra test (JT)[5, 6], in which the test statistic is constructed based on pairwise comparisons between groups, the Cuzick test (CU) [7], which extends the Mann Whitney test (MW) to compare more than two groups, the Terpstra-Magel test [8], which simultaneously compares one observation from each group, and a method more recently proposed by Shan et al. [9], which incorporates the pairwise differences in ranks between observations into the test statistic to capture extra information from the data.

However, existing trend testing methods including rank-based tests are designed to test the null hypothesis that all group means are equal. They do not account for the situation that the group means are unequal but do not follow a monotonic trend. To illustrate, suppose we have data collected on independent samples from three populations. Let the data from the *i*^*th*^ population be denoted by 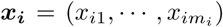, where *m*_*i*_ is the sample size for group *i*. Let *µ*_1_, *µ*_2_ and *µ*_3_ denote the population means of the three groups. The null hypothesis tested by existing methods is *H*_0_ : *µ*_1_ = *µ*_2_ = *µ*_3_, which we refer to as “the narrow null” in which all groups must have the same mean. The alternative hypothesis can be 1-sided such as *H*_*a*_ : *µ*_1_ ≤ *µ*_2_ ≤ *µ*_3_, *µ*_1_ *< µ*_3_ or its 2-sided analogy. Neither the null nor the alternative hypothesis covers what we refer to as a *trendless pattern*, in which the means are unequal but do not follow a monotonic order, for example, *µ*_3_ *< µ*_1_ *< µ*_2_. As a result, while the rejection of the null hypothesis by existing methods indicates that the three-group means are unlikely to be equal, it does not necessarily differentiate a monotonic trend from the trendless pattern. In practice, a trendless pattern is often plausible, in which case it can be useful and important to disentangle a trendless pattern from the truly interesting scenario where the group means do follow a monotonic trend. For example, when a biomarker whose levels in healthy, low-risk cancer and aggressive cancer subjects increase or decrease throughout the three groups, one can use the biomarker to study the prognosis of cancer. However, we argue that existing methods designed for the narrow null are not suited to study such signals because the rejection of the null hypothesis by such methods cannot be interpreted as the presence of a monotonic trend. To the best of our knowledge, no trend testing methods have been published that target the detection and delineation of a monotonic trend from a trendless pattern.

To address this challenge, we propose to formulate the problem using a framework with a less restrictive null hypothesis. We first demonstrate how the application of existing trend testing methods to this problem result in inadequate type 1 error control. We then propose a greedy testing method tailored for the less restrictive null. The greedy test motivated by the likelihood ratio test. Unlike rank-based methods, it does not rely on the assumptions that the shapes and variances of the distributions are identical across groups. The greedy test incurs minimal computational cost and has a simple interpretation related to two-sample t-tests. In Section 4, we use simulation studies to validate the type 1 error control of the method and to evaluate its power, and describe a real data application to illustrate the use of the greedy test. In Section 5, we discuss a more general problem beyond trend testing, in which the proposed test can be used.

## 2 The narrow null versus the broad null

In this paper, we are interested in disentangling a monotonic trend among population means from a trendless pattern where the population means may or may not be equal. Because existing methods ignore the possibility of a trendless pattern in the parameter space, when the *µ*_*i*_’s are unequal but do not follow a monotonic trend, these methods are not guaranteed (and as we will show in the simulation studies in Section 4.1, they often fail) to adequately control type 1 error. To solve this problem, we propose to expand the narrow null hypothesis by adding the trendless pattern to it, thus yielding what we refer to as *the broad null*. In this paper, we consider the case when there are three groups, for which the alternative and the broad null hypotheses can be expressed as:

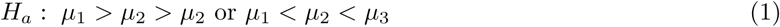

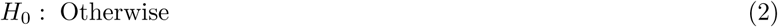

Under this framework, we aim to develop a method that effectively controls type 1 error in the entire null space including the trendless pattern. We denote the parameter space by Θ = {(*µ*_1_, *µ*_2_, *µ*_3_) : *µ*_*i*_ ∈ ℝ, *i* = 1, 2, 3}, and the subspace in Θ corresponding to the alternative hypothesis *H*_*a*_ by Θ_*a*_ = {***µ*** : *µ*_1_ *< µ*_2_ *< µ*_3_ or *µ*_1_ *> µ*_2_ *> µ*_3_} and the subspace corresponding to the broad null *H*_0_ by Θ_0_ = Θ *\* Θ_*a*_. Figure 1 shows the contrasts between the narrow and broad null hypotheses and between their corresponding alternative hypotheses.

**Figure 1:**
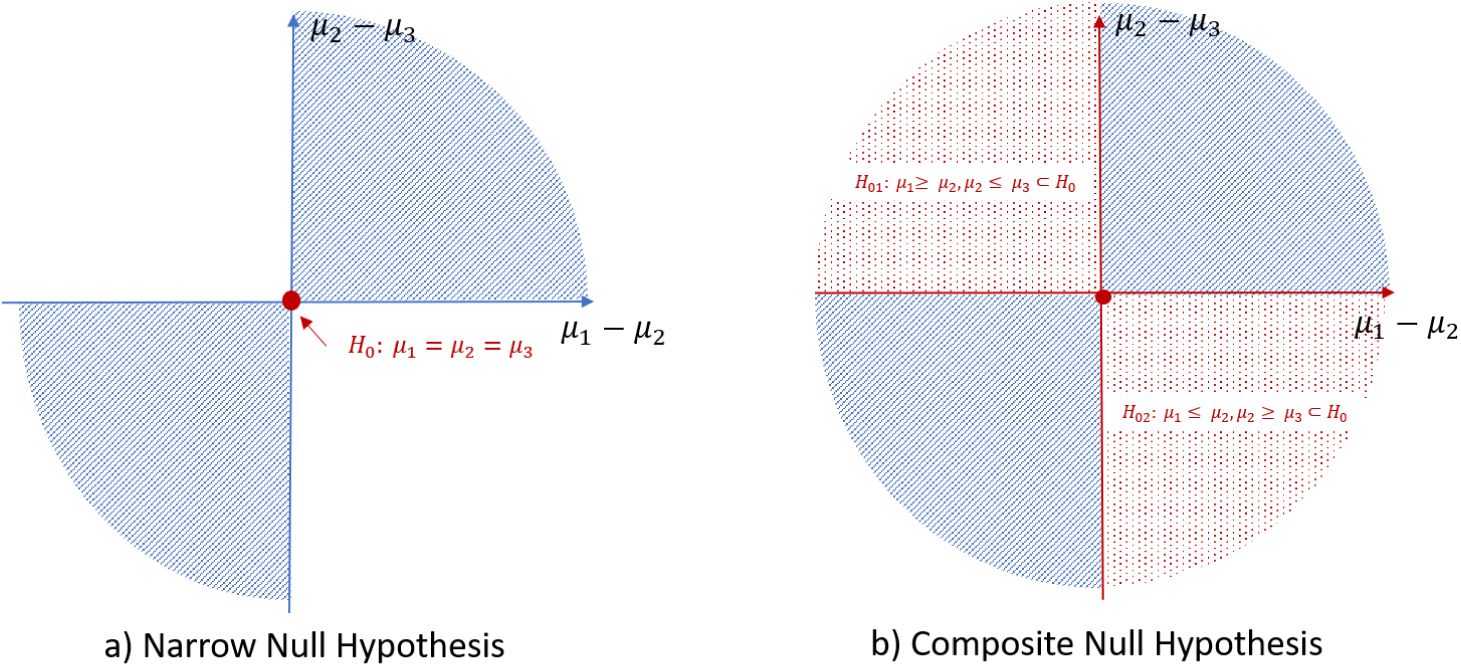
The narrow (Panel a) and broad (Panel b) null hypotheses and the corresponding alternative hypotheses. In each panel, the red region (including areas dotted in red and red solid lines when applicable) represents the null space, and the blue region (including blue solid lines) represents the alternative space.

Although few studies have identified this crucial problem and defined a suitable method to solve it, some naive procedures based two-sample t-tests (TSTTs) may be attempted to tackle it. We will describe one such procedure using 2-sided TSTTs before introducing the greedy test in Section 3.

To utilize TSTTs to test for a strictly monotonic trend in Equation (1), one can first test the hypothesis that *µ*_1_ ≠ *µ*_2_ and *µ*_2_ ≠ *µ*_3_ and then verify that the sample means follow a monotonic trend. For the former, 2-sided TSTTs can be performed comparing groups 1 and 2 and comparing groups 2 and 3. We denote the *p*1 and *p*2 as the p-values of two TSTTs respectively. Because two TSTTs are conducted, the significance level for each test needs to be adjusted. Using Bonferronni correction, for example, one can conduct each TSTT at level *α/*2 for a desired overall type 1 error rate controlled below *α*. If both tests yield significant results at level *α/*2, then both of the two pairs of means are considered significantly different. To conclude that the *µ*_*i*_’s follow a monotonic trend, one can then examine if the 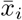’s are monotonic. Namely, if the p-value of this test procedure *p* = 2 ∗ *max*(*p*1, *p*2) is less than *α* and 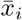’s follow a monotonic trend, we can reject the null hypothesis. In table 1, we detail the specific steps in this procedure, which we will refer to as the naive procedure based on 2-sided TSTTs.

**Table 1:** Naive testing procedure based on 2-sided TSTTs.

| Testing Steps | Test hypothesis |
| --- | --- |
| Step 1.<br>Compare $\mu_1$ and $\mu_2$ | Test 1 (level $\alpha/2$):<br>$H_{01} : \mu_1 = \mu_2$<br>$H_{a1} : \mu_1 \neq \mu_2$ |
| Step 2.<br>Compare $\mu_2$ and $\mu_3$ | Test 2 (level $\alpha/2$):<br>$H_{02} : \mu_2 = \mu_3$<br>$H_{a2} : \mu_2 \neq \mu_3$ |
| Step 3.<br>Check the order of the sample means | $\bar{x}_1 > \bar{x}_2 > \bar{x}_3$<br>or<br>$\bar{x}_1 < \bar{x}_2 < \bar{x}_3$ |
| Reject the overall $H_0$ if both of the following hold:<br>(i) both tests 1 and 2 yield a rejection, and (ii) the sample means have a monotonic order | |

Two other naive procedures are described in Section 5. We additionally note that the Delta method is not applicable for this problem. To attempt the Delta method, one would consider *η* = *λ*_1_ *·λ*_2_ to be the parameter of interest, where *λ*_1_ = *µ*_1_ − *µ*_2_ and *λ*_2_ = *µ*_2_ − *µ*_3_, and construct a test based on 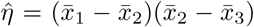. However, the conditions for the multivariate Delta method do not hold because the gradient of 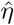 is zero when *λ*1 = *λ*_2_ = 0[10].

## 3 A trend test for the broad null

We focus on the problem of testing the hypothesis in Equation (1). We allow the sample sizes and the population variances to be different between the three groups. We assume a reasonably large sample size for all three groups so that the sample mean of each group is well approximated by a normal distribution based on the central limit theorem. The asymptotic distributions of the sample means are given by 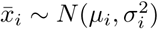, where *µ*_*i*_ is the population mean of group *i* and 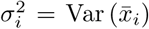 is the variance of the *i*^*th*^ sample mean for *i* = 1, 2, 3.

### 3.1 A test statistic motivated by the likelihood-ratio test

We will first tackle the case if *σ*_*i*_’s are known, and the general case with unknown *σ*_*i*_’s will be discussed later (Section 3.2). Ignoring constants that do not depend on the parameters *µ*_*i*_’s, the log-likelihood function can be written as 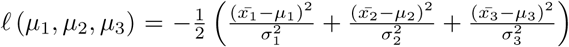. To derive the likelihood ratio test, we start by finding the restricted and unrestricted maximum likelihood estimators (MLEs) for the unknown parameters *µ*_1_, *µ*_2_ and *µ*_3_. It is easily seen that the unrestricted MLE is given by the sample means 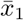, 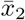 and 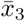. The restricted MLE which maximizes the likelihood function in the null space, we need to consider two situations, depending on whether 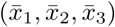 is consistent with Θ_0_. We therefore divide the “data space” 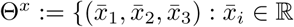, *i* = 1, 2, 3} into two parts (Figure 2), one consistent with Θ_*a*_ and one with Θ_0_:

**Figure 2:**
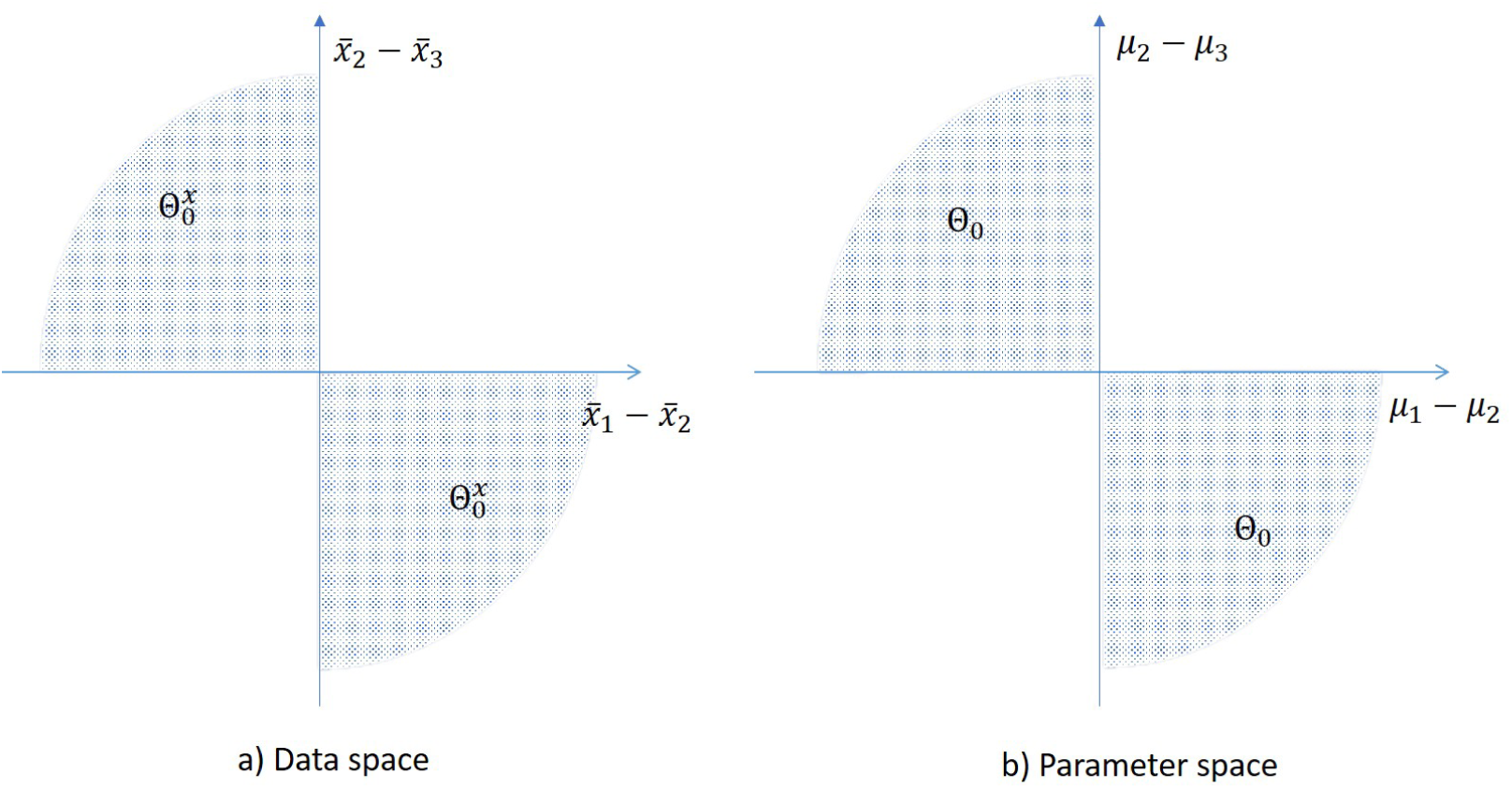
Partitions of the data space Θ^*x*^ and the parameter space Θ.

**Figure 3:**
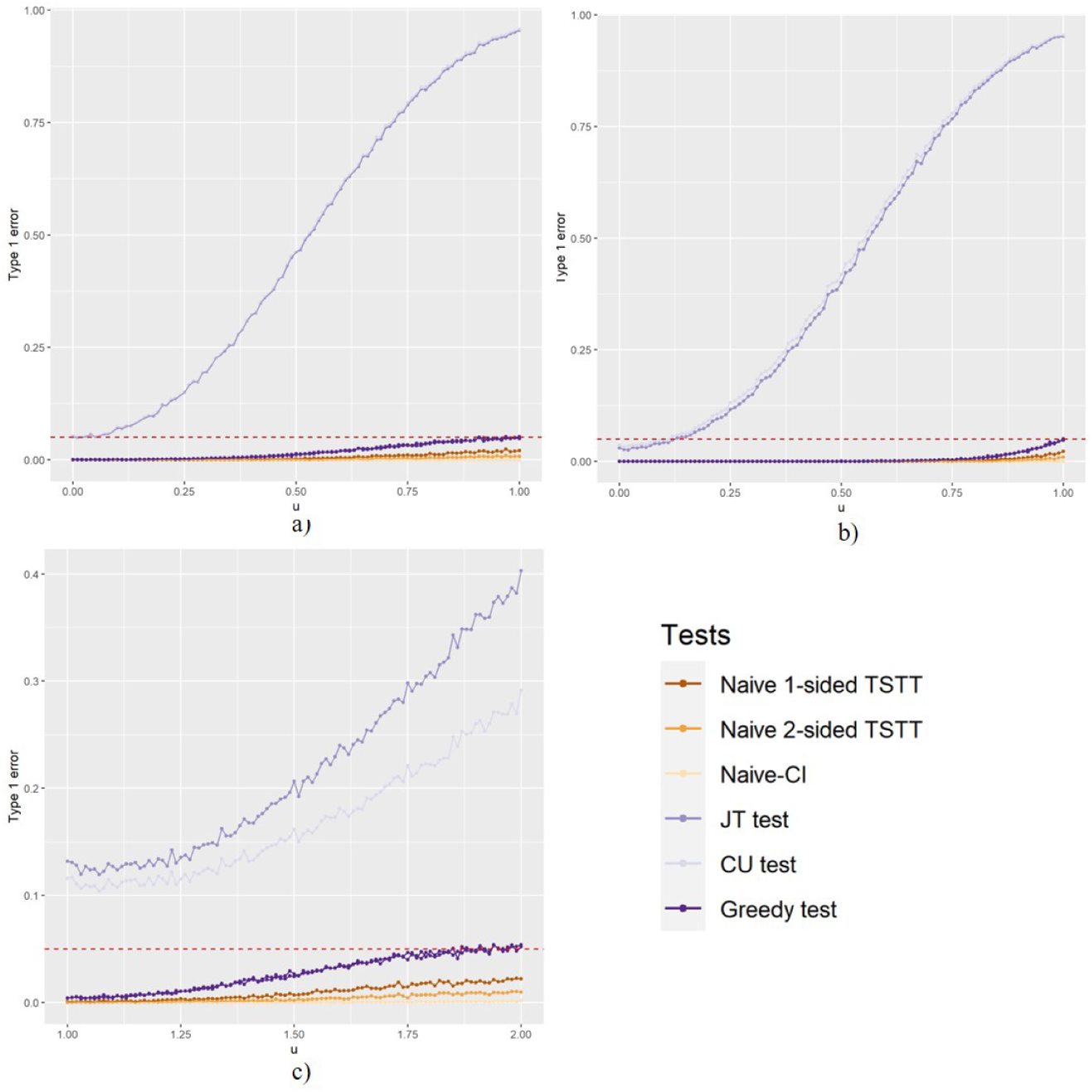
Empirical type 1 error results. The three panels compare the empirical type 1 error rates of the greedy test, CU, JT and three naive procedures under settings I) boundary of null, II) interior of null and III) unequal variances and different distribution families, respectively. Each curve shows the empirical type 1 error rate (vertical axis) as a function of *mu*_3_ (horizontal axis). The curves are color coded to represent the methods in comparison. The horizontal dashed line display the nominal level at 0.05. The greedy test and the naive procedures consistently controls type 1 error effectively, whereas both CU (grey curve) and JT (light purple curve) fail to control type 1 error in the majority of the settings.

**Figure 4:**
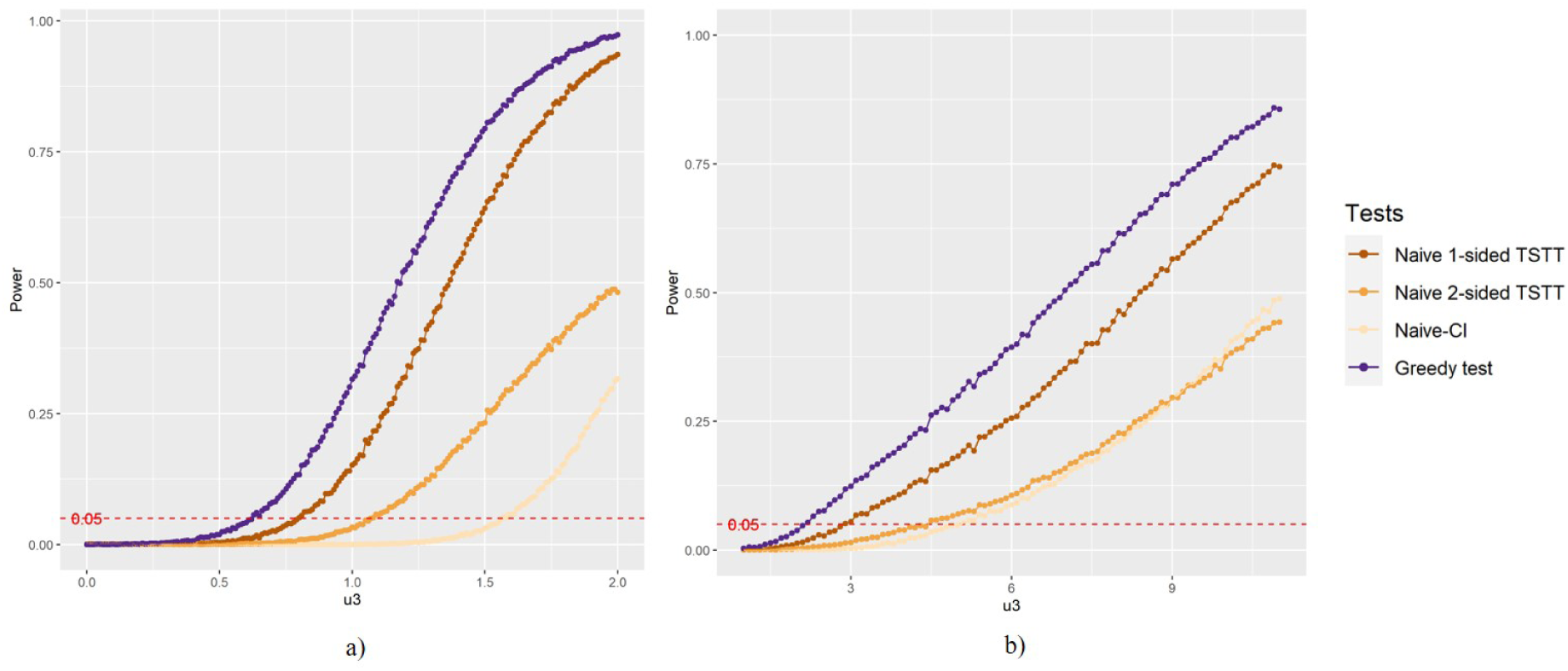
Empirical power results of the greedy test and three naive procedures for settings IV and V. Each curve shows the empirical power (vertical axis) of a method as a function of *µ*_3_ (horizontal axis). The curves are color coded to represent the four methods. Across the settings, the greedy test (purple curve) is the most powerful, followed by the naive procedure based on 1-sided TSTTs (Brown curve), the naive procedure based on 2-sided TSTTs (dark yellow curve), and the naive procedured based on confidence intervals (light yellow curve).

**Figure 5:**
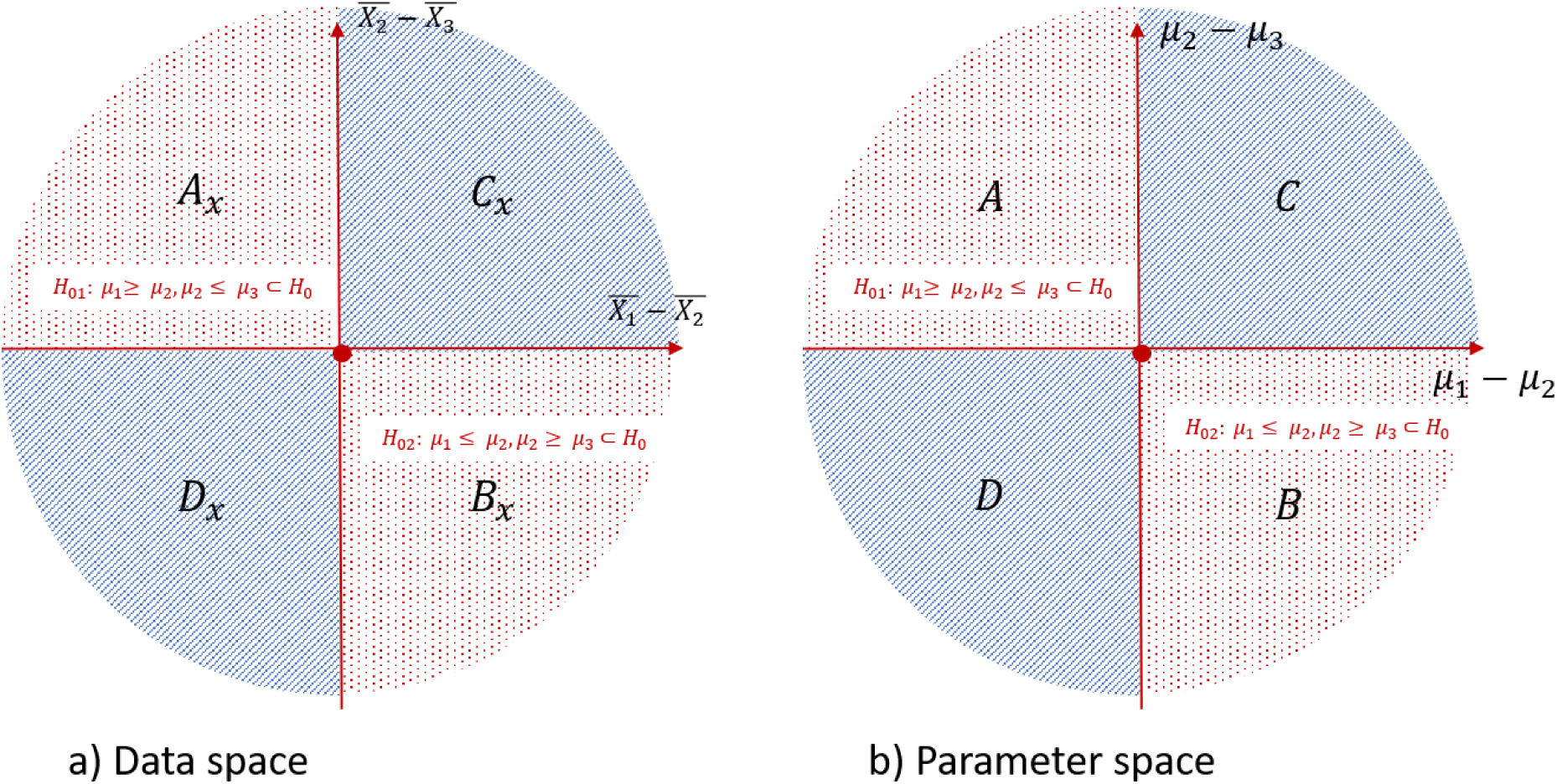
Further partitions of the data space Θ^*x*^ and the parameter space Θ.

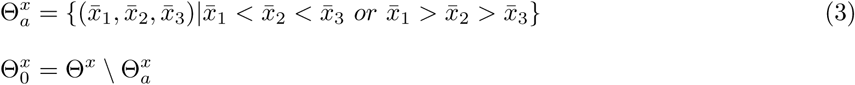

The following lemma concerns the restricted MLE.

**Lemma 1.**

1. *When* 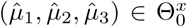, *the restricted MLE which maximizes the log-likelihood function over* Θ_0_ *is* 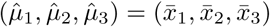.
2. *When* 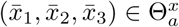, *the restricted MLE which maximizes the log-likelihood function over* Θ_0_ *is*

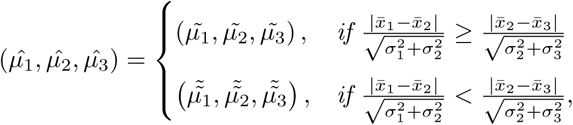

*where* 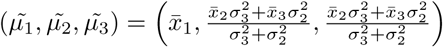, 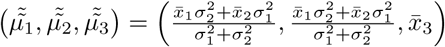.

To intuitively explain the Lemma, when 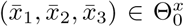, the data are consistent with the pattern in the null hypothesis and therefore the restricted MLE coincides with the unrestricted MLE.

When 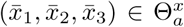, under the null, the MLEs are attained when two populations share the same mean. In other words, the MLEs are taken on one of the two boundaries of Θ_0_, which is not surprising given that the trend in the data does not align with the ordering of the population means. The smaller of the two standardized distances (i.e. 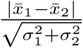 and 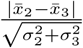 reveals which of the two boundaries of Θ_0_ yields a greater log likelihood.

Given the restricted and unrestricted MLEs, we can derive the maximum value of the log-likelihood function under restricted and unrestricted cases and hence the likelihood ratio. First, we derive the maximum of the log-likelihood under Θ_0_. When 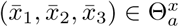, plugging in the MLEs for (*µ*_1_, *µ*_2_, *µ*_3_) given in Lemma 1 into (*µ*_1_, *µ*_2_, *µ*_3_) yields a log-likelihood value that depends on the standardizes distances. When the sample means do not follow a monotonic trend, the restricted MLEs coincide with the unrestricted MLEs, and thus the likelihood ratio statistic is zero. These results are summarized in Theorems 2 and 3.

**Theorem 2.** *When* 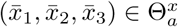, 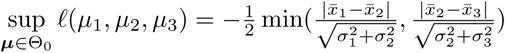

**Theorem 3.** *The likelihood ratio statistics is given by*,

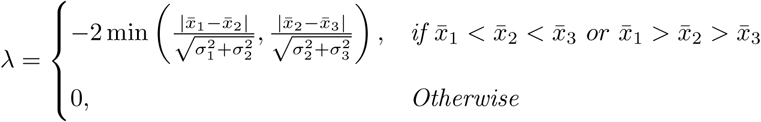

This statistic makes intuitive sense as a way of measuring how inconsistent the data is with *H*_0_, because when sample means have a monotonic trend and the minimal of the distances between sample means is large, the population means are likely to have a strictly monotonic trend.

Motivated by the form of the likelihood ratio test, we define our test statistic as follows

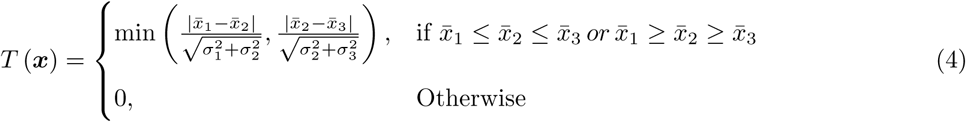

### 3.2 Assessing Statistical Significance

Let *t* be the realized value of the test statistic defined in Equation (2). For fixed ***µ***, *P*_***µ***_(*T* (***x***) *> t*) is a measure of compatibility of the observed data with null hypothesis. To effectively control type 1 error in the entire null space, we obtain the p-value as 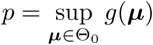. With some reparameterization, we will show that the p-value has a simple and intuitive analytical form.

Recall that *λ*_1_ = *µ*_1_ − *µ*_2_ and *λ*_2_ = *µ*_2_ − *µ*_3_. Note that *H*_0_ can be equivalently expressed in terms of *λ*_1_ and *λ*_2_ as *λ*_1_*λ*_2_ *>* 0. It is not hard to show that *P*_***µ***_(*T* (***x***) *> t*) depends on ***µ*** only through *λ*_1_ and *λ*_2_. We hence let 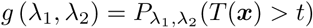, and it follows that the p-value is given by 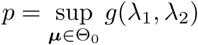. Under the null, the population means have higher chance to yield strictly monotonically ordered sample means when two of the sample means are equal. Therefore, the measure of compatibility is higher when one of the *λ*s is zero and lower when otherwise. In fact, the measure of compatibility is maximized when one of the *λ*’s is zero and the other approaches infinity. This is formalized in the following Lemma.

**Lemma 4.** 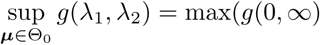 = max(*g*(0, ∞), *g*(∞, 0), *g*(0, −∞), *g*(−∞, 0))

By evaluating the right hand side of the equation above, the following Theorem for the p-value follows.

**Theorem 5.** *The p-value is given by* 1 − Φ(*t*), *where* Φ(*·*) *is the cumulative distribution function of N* (0, 1) *and t is the observed value of the test statistic T*.

With the great simplicity of the form for the p-value, the greedy test can be implemented using a two-step procedure: 1) find the test statistic by taking the minimum of the two standardized distances. 2) derive the p-value by obtaining the upper tail cumulative probability for the standard normal distribution. It is noteworthy that the test only depends on the standardized distances between the group means and is unaffected by the amount of correlation between the standardized distances.

In practice, when 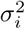’s are unknown, we replace them with a consistent estimator such as the sample variance estimator given by 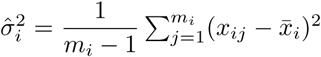 in the test statistic defined in Equation (4). The test statistic hence becomes

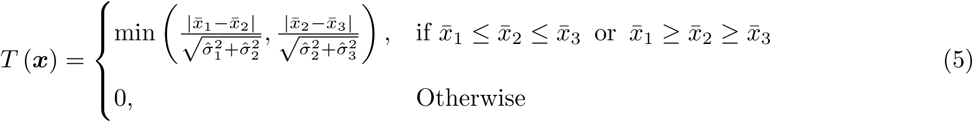

### 3.3 Rejection region and connection with two-sample t-tests

Given a desired significance level *α*, it is easily seen that the rejection region of the greedy test is given by {***x*** : *p < α*} = {***x*** : *T*_1_(***x***) *· T*_2_(***x***) *>* 0, |*T*_1_(***x***)| *> z*_1−*α*_, |*T*_2_(***x***)| *> z*_1−*α*_}, where 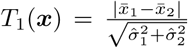 and 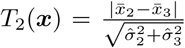 are 2-sided TSTT statistics, and *z*_1−*α*_ is the 1 − *α* quantile of the standard normal distribution. This brings about a natural interpretation of the greedy test method based on TSTTs. It can be equivalently implemented using the following simple procedure:

1. Are the sample means located in 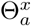 (i.e. do the sample means follow a monotonic trend)? If so, go to Step 2; If not, fail to reject the null (p-value *>* 0.5).
2. Conduct 2-sided TSTTs at level 2*α* between groups 1 and 2 and between groups 2 and 3.
3. Follow either one of the follow two procedures, which are equivalent:
  - Reject the null if both TSTTs reject their corresponding nulls at level 2*α*.
  - Equivalently, let the p-values of the TSTT’s be *p*_1_ and *p*_2_. The final p-value of the greedy test can be obtained by taking *p* = max(*p*_1_, *p*_2_)*/*2. Reject the null if *p < α*.

It is worth noting that the procedure described above is closely connected to the naive procedure based on 2-sided TSTTs (see Section 2 and Table 1). The key difference between the two is that the greedy test requires both 2-sided TSTTs to be performed at level 2*α*, whereas the naive procedure performs both TSTTs at level *α/*2.

In other words, the p-values for 2-sided TSTT is a linear transformation from the greedy test. Recall the p-value for 2-sided TSTT is *p*_2−*sidedT ST T*_ = 2 max(*p*_1_, *p*_2_), so *p*_2−*sidedT ST T*_ = 4*p*_*greedy*_. Therefore, the greedy test is expected to achieve substantial power gain over the 2-sided TSTT procedure and the listed naive procedures in Appendix C.

## 4 Results

### 4.1 Simulation studies

First, we verify the effective type 1 error control of the greedy test and three naive procedures: one based on 2-sided TSTTs as described in Section 2, one based on 1-sided TSTTs and one based on confidence intervals (see Appendix C). In addition, we show that existing methods proposed for the narrow null do not control type 1 error effectively. As examples of rank-based methods, we focus on JT and CU, which, like other rank-based tests, are proposed to test the narrow null and further assume that the populations follow the same distribution under the null.

We consider the following simulation settings from the broad null.

I. Boundary of the null: Two of the groups have the same mean, which differs from the mean of the third group. The three groups are assumed to follow N(0,1), N(0,1), and N(*µ*_3_, 1), respectively, where *µ*_3_ is varied from 0 to 1.
II. Interior of the null: All three group means are different but do not follow a monotonic trend. The three groups are assumed to follow N(0,1), N(1,1), and N(*µ*_3_, 1), respectively, where *µ*_3_ is varied from 0 to 1.
III. Unequal variances and different distribution families: The three groups do not come from the same distribution family and do not have the same variance. The three groups are assumed to follow N(1,10), N(1,1), and Poisson(*µ*_3_), respectively, where *µ*_3_ is varied from 1 to 2.

For each simulation setting, we simulate *m*_*i*_ = 30 samples independently from each of the three populations. Then, the type error 1 rate of a method is estimated based on 10,000 replicates as the frequency at which the p-value is below a nominal level of *α* = 0.05.

Across the simulation settings, the greedy test as well as the naive methods effectively control type 1 error at level 0.05. However, both CU and JT suffer from severely inflated type 1 error for a vast majority of the settings. Settings I and II assume normally distributed data with equal variances across groups. In setting I (Figure 1a), the first two groups have the same mean and hence the parameters are located on the boundary of the broad null which we aim to test. In this setting, the parameters are consistent with the narrow null if and only if *µ*_3_ = 0, in which case type 1 error is also well controlled by the rank-based tests. However, when *µ*_3_ *>* 0, the parameters are treated as being part of the alternative space by rank-based methods. As such, the rejection probability tends to fall well above the nominal level 0.05 for JT and CU.

In setting II, *µ*_1_ ≤ *µ*_3_ ≤ *µ*_2_. When *µ*_3_ is strictly between *µ*_1_ and *µ*_2_, the group means are all unequal but not monotonic. This setting is in fact treated by rank-based methods as implausible because it belongs neither to the narrow null nor to the alternative hypothesis targeted by rank-based methods. In this case, it is not surprising that rank-based methods do not guarantee adequate control of type 1 error. In fact, JT and CU can result in excessive false positives with a rate of up to 100%. In summary, rank-based methods are severely mis-calibrated both on the boundary and in the interior of the broad null hypothesis, whereas the greedy test and the naive procedures effectively control type 1 error globally.

For setting III, the three populations have distributions from different families. When *µ*_3_ *>* 1, CU and JT result in excessive type 1 error for the same reason that they do in Setting I. In contrast to Setting II, the three groups follow distributions from different families and have unequal variances, which is a violation of the assumption of rank-based methods that the groups have identical distributions under the null. This explains why CU and JT have inflated type 1 error rate even when *µ*_1_ = *µ*_2_ = *µ*_3_ = 1. Effective type 1 error control by the greedy test and the naive procedures, however, do not rely on the identical distribution assumption.

We then evaluate the power of our approach and compare it with the naive methods. Rank-based methods are not included in the power simulations, because they do not control type 1 error adequately. To this end, we consider two settings under the alternative hypothesis.

IV. Increasing trend: The three groups follow normal distributions with monotonically increasing means and the same variance. The three groups are assumed to follow *N* (0, 1), 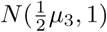, and *N* (*µ*_3_, 1), respectively, where *µ*_3_ is varied from 0 to 1.
V. Unequal variances and different distributions families: The three groups do not come from the same distribution family and do have the same variance. The group means follow an increasing trend. The three groups are assumed to follow *N* (1, 10), 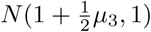, and Poisson(1 + *µ*_3_), respectively, where *µ*_3_ is varied from 0 to 10.

In both settings, power grows for all methods as *µ*_3_ increases. Throughout the simulations, the greedy test achieves the highest power. This demonstrates the power advantage of our method over the naive procedures.

### 4.2 Real data application

We illustrate the proposed test by applying it to the gut microbiome data produced during a prior investigation of the zebrafish gut microbiome Gaulke et al. 2019 [11]. This prior study aimed to investigate the link between the gut microbiome and gastrointestinal parasite infection and pathology. As part of the study, 105 zebrafish were exposed to *Pseudocapillaria tomentosa* or controls, a helminthic parasite that infects the zebrafish gut to induce pathological changes including intestinal inflammation, tissue damage, and hyperplasia Kent. The study profiled the composition of the gut microbiome through 16S rRNA gene sequence analysis, quantified total worm burden through cytology, and measured the extent of histopathological damage induced by the infection.

Specifically, the gut microbiome of the fish was measured over 12 weeks of infection via cross-sectional 16S sequencing, which generated the relative abundances of 565 genera for each fish. Two types of pathological changes on the parasite-exposed fish, inflammation and hyperplasia, were assessed by a pathologist based on examination of intestine tissues. For either type of changes, severity was scored between 0 and 3 for each fish. In the prior work, the investigators modeled how the variation in the relative abundance of specific taxa explained the variation in histopathological outcomes, but did not resolve monotonic relationships between these features *per se*.

We use these data in conjunction with our application to resolve microbiota that specifically monotonically associate with histopathological outcomes, which are useful to detect because such associations point to intimate relationships between pathology and the taxon in question. Such taxa are consequently strong candidates for being functionally intertwined with the pathological phenotype. To detect such taxa, we group the samples into three categories based on the inflammation score: no inflammation (score = 0, n = 47), mild to moderate inflammation (score = 1 or 2, n = 45), and severe inflammation (score = 3, n = 10). We also group the samples based on the hyperplasia score: no hyperplasia (score =0, n = 32), mild and moderate hyperplasia (score = 1 or 2, n = 53), and severe hyperplasia (score = 3, n = 17).

We first aim to detect overall patterns of the gut microbiome that associate with inflammation severity. To this end, we perform non-metric multidimensional scaling (NMDS) based on Bray-Curtis dissimilarity and principal component analysis (PCA), on the microbiome data. For either NMDS or PCA, we retain the top three components, which are then tested individually for monotonic trends with inflammation severity. Bonferroni-corrected p-values are reported in Table 2 based on the proposed greedy test, the naïve procedure based on 1-sided TSTT, and the naïve procedure based on 2-sided TSTT. The greedy test finds strong evidence that the second NMDS component varies strictly monotonically across the severity groups of inflammation (MDS2, Bonferroni corrected p = 0.0030). The monotonic trend is also identified by the two naïve procedures, although with less significant p-values (Bonferroni-corrected p = 0.0059 and 0.012, respectively). Based upon the PCA results, the greedy test identifies a statistically significant monotonic trend associated with the second principal component (PC2, Bonferroni-corrected p = 0.037). The naïve procedures (Bonferroni-corrected p = 0.074 and 0.15 respectively), however, fail to capture this signal at level 0.05 due to their lower power compared with the greedy test. These results indicate that there exist overall features of the gut microbiome that vary monotonically with increasing severity level of inflammation in the intestine among parasite-exposed zebrafish.

**Table 2:** Bonferroni-corrected p-values in zebrafish gut microbiome analysis, based on the greedy test, the naive procedure based on 1-sided TSTT (Naive 1-sided) and the naive procedure based on 2-sided TSTT (Naive 2-sided). P-values below 0.05 are in bold type, and p-values between 0.05 and 0.2 are italicized. Microbial features for which none of the methods yields a raw p-value below 0.05 are omitted.

| NMDS |  |  |  |
| --- | --- | --- | --- |
|  | Greedy test | Naive 1-sided | Naive 2-sided |
| MDS2 | **0.0030** | **0.0059** | **0.012** |
| PCA |  |  |  |
|  | Greedy test | Naive 1-sided | Naive 2-sided |
| PC2 | **0.037** | 0.074 | 0.15 |
| Genus-level Tests |  |  |  |
|  | Greedy test | Naive 1-sided | Naive 2-sided |
| *Cetobacterium* | 0.11 | 0.22 | 0.44 |

**Table 3:** Naive procedure based on 1-sided TSTTs

|  |  |
| --- | --- |
| Step 1. Compare $\mu_1$ and $\mu_2$ | Test 1 (level $\alpha/2$):<br>$H_{01} : \mu_1 \geq \mu_2$<br>$H_{a1} : \mu_1 < \mu_2$ |
| Step 2. Compare $\mu_2$ and $\mu_3$ | Test 2 (level $\alpha/2$):<br>$H_{02} : \mu_2 \geq \mu_3$<br>$H_{a2} : \mu_2 < \mu_3$ |
| Step 3. Compare $\mu_1$ and $\mu_2$ | Test 3 (level $\alpha/2$):<br>$H_{03} : \mu_1 \leq \mu_2$<br>$H_{a3} : \mu_1 > \mu_2$ |
| Step 4. Compare $\mu_1$ and $\mu_2$ | Test 4 (level $\alpha/2$):<br>$H_{04} : \mu_2 \leq \mu_3$<br>$H_{a4} : \mu_2 > \mu_3$ |
| Reject the overall $H_0$ if one of the following happens:<br>(i) both tests 1 and 2 yield a rejection, or<br>(ii) both tests 3 and 4 yield a rejection | |

Building upon these results, we next apply our procedure to resolve specific taxa that monotonically associate with inflammation severity. We focus the analysis on the common genera which are present in at least 20% of the samples. A total of 49 such genera are identified in the data. The relative abundance of each genus is tested for monotonic trends across inflammation severity groups using the greedy test and two naïve procedures (Table 2). Based on the greedy test, there is suggestive evidence that *Cetobacterium* decreases monotonically as inflammation severity elevates from no inflammation to severe inflammation (Bonferroni-corrected p = 0.11, 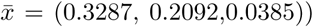. This finding reinforces the discovery of a negative (and not necessarily monotonic) association between Cetobracterium and parasite burden in Gaulke et al, 2019. We note that, using the naïve trend testing procedures, the monotonic trend associated with *Cetabacterium* would yield substantially greater p-values (Bonferroni-corrected p = 0.22 and 0.44, respectively).

Similar analyses are performed for the groups defined based on hyperplasia scores, and no significant signals are found. Overall, our results indicate that while both the greedy test and the naïve procedures do well in capturing highly significant monotonic trends, the power gain achieved by the greedy test has the potential of enabling the discoveries of mild to moderate signals that would be missed by the use of the naïve tests.

## 5 Discussion

We have proposed a greedy testing method for a strictly monotonic trend among the means of three independent groups against a composite null hypothesis that the group means are equal or unequal but are not strictly monotonic. We demonstrate both theoretically and empirically that existing testing methods are not valid for this problem because they target a narrow null in which the group mean are all equal. To address this issue, we have developed a greedy testing method that achieves global type 1 error control in the entire null space. Moreover, unlike most nonparametric methods based on rank, the proposed method does not require the groups to have the same distribution under the null or the same variance and therefore is more flexible. The greedy test is computationally simple and the test statistic has a simple expression because of its connection with TSTTs. However, compared to a naive procedure based on pairwise TSTTs, the greedy test improves power substantially because we have shown that, for each pairwise comparison, it only requires a significance level that is four times the significance level entailed by the naive procedure to achieve global type 1 error control.

We have shown in simulation studies that methods intended for testing the narrow null result in excessive false positives when used to test the broad null, whereas the greedy test effectively controls the type 1 error rate in a wide range of settings including those in which the groups have distributions from different families and different variances. In addition, we have demonstrated that the proposed test outperforms three naive procedures that we have described in power. Finally, we have applied the method to the data from GSEP, which is a case-control study on prostate cancer. We have found suggestive evidence that Gleason score increases consistently across the three age groups defined as (age *<* 60, 60 ≤ age ≤ 70,age *>* 70) and that PSA level increases consistently as the minor allele count for rs851023 goes from 0 to 2.

Although the development of the method has been primarily motivated by the trend test problem with a broad null, the method can be used to solve a more general class of problems beyond trend tests. To see this, we first write the hypotheses in (1) and (2) in the following form

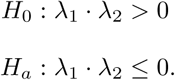

and write the test statistic in Equation (4) as

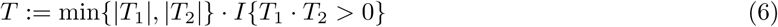

In the context of a trend test between three groups, *λ*_1_ and *λ*_2_ represent the pairwise differences in means and *T*_1_ and *T*_2_ are the corresponding 1-sided test statistics. However, the proof of Lemma 4 does not rely on these specific definitions of *λ*_1_ and *λ*_2_. In a more general framework, *λ*_1_ and *λ*_2_ can be any parameters for which it is of interest to test whether they are both nonzero and have the same sign. For this general setup, the test statistic defined in Equation (6) can be used and its p-value can be assessed using the expression given in Theorem 5. The proof of Lemma 4 in the Appendix is still valid to show that Theorem 5 holds in the more general setup, so long as the following conditions hold: a) *T*_1_ and *T*_2_ are valid 2-sided statistics for *λ*_1_ = 0 and *λ*_2_ = 0, respectively; b) (*T*_1_, *T*_2_) has an (asymptotic) bivariate normal distribution, *E*(*T*_*i*_) has the same sign as *λ*_*i*_, var(*T*_*i*_) = 1, and cov(*T*_1_, *T*_2_) ≤ 0. In particular, this includes the setting in which *T*_1_ and *T*_2_ are independent z-test statistics. This is potentially relevant to many medical applications. For example, when two separate studies both evaluate the association between a genetic factor and a disease, the test statistics are independent provided that the studies are conducted on separate cohorts. In this case, the greedy test has the potential to take advantage of combining the summary statistics of the two studies to boost power by enabling the detection of a significant association at level *α* so long as the p-values from both studies are below 2*α*.

## Acknowledgment

I would like to express my gratitude to Professor Duo Jiang for extraordinary support and encouragement for this work, for commenting on earlier drafts of this paper, and for advising on the presentation and positioning of this paper. I also would like to thank Professor Yuan Jiang and Professor Thomas Sharpton for their valuable feedback.

## Appendix A

In this Appendix, we show the results in Section 3.1

### A1. Proof of Lemma 1

*Proof.* We start by introducing some notation that we will use to partition the parameter space Θ and the data space 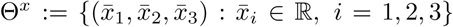. First, we define the four disjoint regions of the parameter space illustrated in Figure 1b as follows (see Figure 5a)

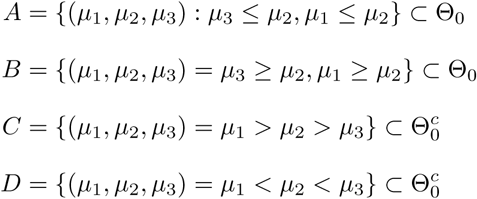

Analogous regions in the data space can also be defined (see Figure 5b):

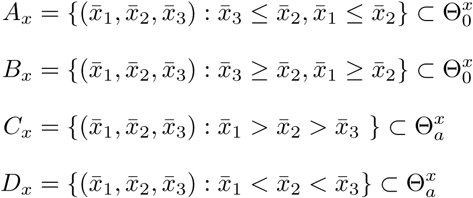

We will present the proof of Lemma 1 for the case when 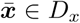. For 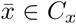, the results can be shown using a similar argument. Assuming 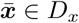, we first prove the following lemma on the maximization of the log-likelihood function *l* (*µ*_1_, *µ*_2_, *µ*_3_) for ***µ*** ∈ *A*.

#### Lemma 6.

*Assuming* 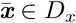, *we have*

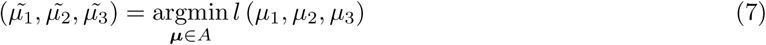

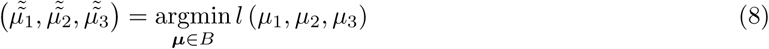

*where* 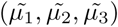 *and* 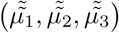 *are as defined in Lemma 1.*

*Proof.* We first note that 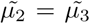 and 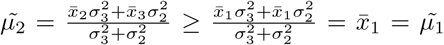, where the last inequality holds because 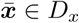. This implies that 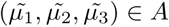. To show equation (7), it remains to show that

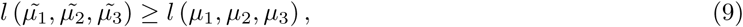

for any (*µ*_1_, *µ*_2_, *µ*_3_) ∈ *A*. To this end, we note that for such (*µ*_1_, *µ*_2_, *µ*_3_) ∈ *A*,

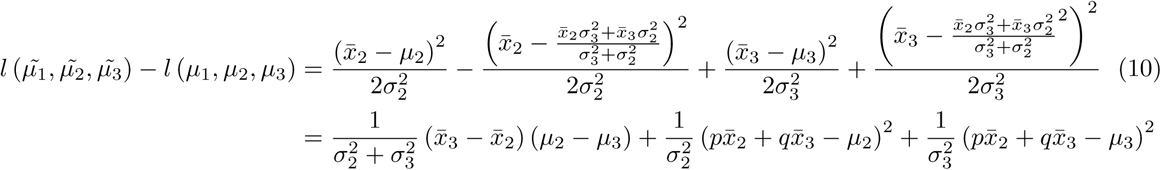

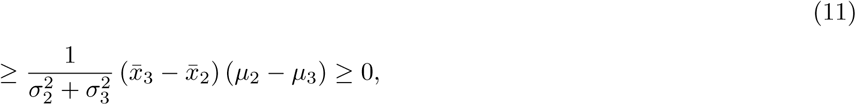

where 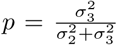, 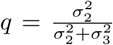 and the last inequality follows from *µ*_2_ ≥ *µ*_3_ and 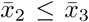. Equation (7) is thereby proved. Equation (8) can be shown using a similar argument.

The following lemma evaluates the log likelihood function at the optima given in Lemma 6.

#### Lemma 7.

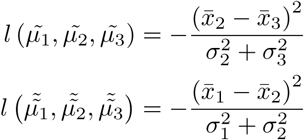

The proof of Lemma 7 is a straightforward substitution into the log likelihood function and is omitted here.

Combing Lemma 6, Lemma 7, and Θ_0_ = *A* ∪ *B*,

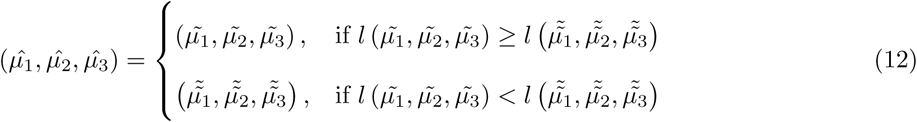

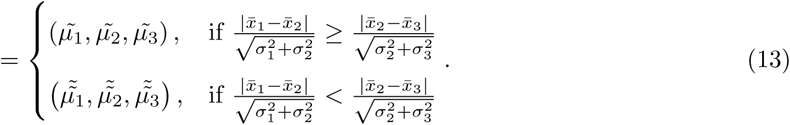

Lemma 1 is thereby proved.

### A2. Proof of Theorem 2

*Proof.* Combining Lemma 1, Lemma 6, and Lemma 7 yields

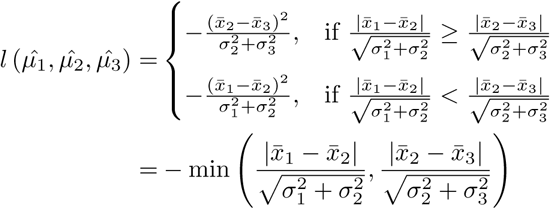

## Appendix B

In this appendix, we prove the results in Section 3.2.

### B1. Proof of Lemma 4 and Theorem 5

*Proof.* To show Lemma 4 and Theorem 5, we first show that *g*(0, ∞) = *g*(∞, 0) = *g*(0, −∞) = *g*(−∞, 0) = 1 − Φ(*t*). Recall that 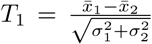 and 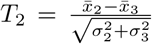. Notice that *T* and *T* have a bivariate normal distribution:

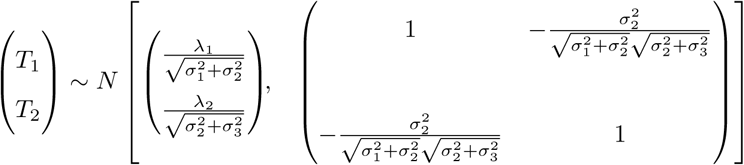

Without loss of generality (WLOG), we focus on the case when 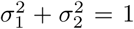 and 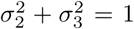 for this proof, so

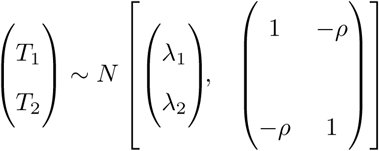

where *ρ* = −corr(*T*_1_, *T*_2_) and *ρ >* 0.

We shift the center of the bivariate normal distribution to (0, 0), and let *Z*_1_ = *T*_1_ − *λ*_1_ and *Z*_2_ = *T*_2_ − *λ*_2_. Then *Z*_1_ and *Z*_2_ have a bivariate normal distribution

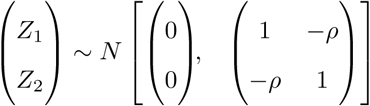

We can now express *g*(*λ*_1_, *λ*_2_) in terms of *Z*_1_ and *Z*_2_ as

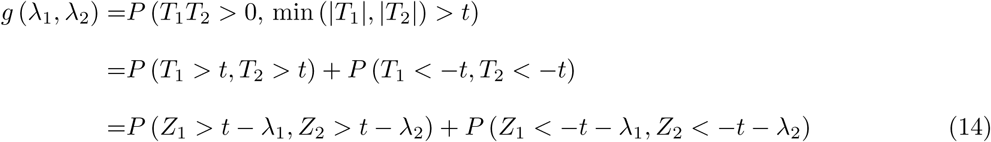

Hence, 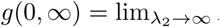 *P* (*Z*_1_ *> t, Z*_2_ *> t* − *λ*_2_) + 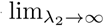 *P* (*Z*_1_ *<* −*t, Z*_2_ *<* −*t* − *λ*_2_) = *P* (*Z*_1_ *> t*) = 1 − Φ(*t*). Likewise, it can be shown that *g*(∞, 0) = *g*(0, −∞) = *g*(−∞, 0) = 1 − Φ(*t*).

To show Lemma 4 and Theorem 5, it remains to show that *g*(*λ*_1_, *λ*_2_) ≤ *P* (*Z > t*) for *λ*_1_, *λ*_2_ ∈ Θ_0_, where *Z* is a standard normal random variable. In equation (14), it is apparent that the magnitudes of *λ*_1_ and *λ*_2_ directly affect the value of *g*(*λ*_1_, *λ*_2_). In fact, *ρ* also plays a role in *g*(*λ*_1_, *λ*_2_) as it influences the distribution of (*Z*_1_, *Z*_2_). To highlight the dependence on *ρ*, for rest of the proof, we write *g*_*ρ*_(*λ*_1_, *λ*_2_). We introduce Lemma 8 which states that for all *λ*_1_, *λ*_2_ ∈ Θ_0_ and when *corr*(*T*_1_, *T*_2_) ≤ 0 (i.e. *ρ <* 0), *g*_*ρ*_(*λ*_1_, *λ*_2_) is maximized when *ρ* = 0. With this lemma, it suffices to show that Lemma 4 holds when *ρ* = 0.

#### Lemma 8.

*g*_*ρ*_(*λ*_1_, *λ*_2_) ≤ *g*_*ρ*=0_(*λ*_1_, *λ*_2_).

The proof of Lemma 8 will be provided in Appendix B2.

With this lemma, it suffices to show that Lemma 4 holds when *ρ* = 0. WLOG, we focus on the case when (*λ*_1_, *λ*_2_) ∈ *B* and |*λ*_1_| *>* |*λ*_2_|. Let *a* = −*λ*_2_. Then, 0 *< a < λ*_1_, and *g*(*λ*_1_, *λ*_2_) = *P* (*Z*_1_ *> t* − *λ*_1_, *Z*_2_ *> t* + *a*)+ *P* (*Z*_1_ *> t* + *λ*_1_, *Z*_2_ *<* −*t* + *a*). When *ρ* = 0, *Z*_1_ and *Z*_2_ are independent. Thus, *g*_*ρ*=0_(*λ*_1_, *λ*_2_) = *P* (*Z*_1_ *> t* − *λ*_1_)*P* (*Z*_2_ *> t* + *a*) + *P* (*Z*_1_ *<* −*t* − *λ*_1_)*P* (*Z*_2_ *<* −*t* + *a*). To show *g*_*ρ*=0_(*λ*_1_, *λ*_2_) ≤ 1 − Φ(*t*) = *P* (*Z > t*), we will discuss three cases depending on the magnitude of *t*: Case A. 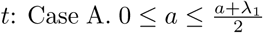, Case B. 0 ≤ *t* ≤ *a* ≤ *λ*_1_, and Case 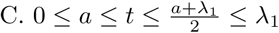.

*Case A* When 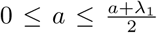 < *t*, we know that *t* − *λ*_1_ *>* −*t* + *a*. Therefore, *P* (*Z*_1_ *> t* − *λ*_1_) + *P* (*Z*_1_ *<* −*t* + *a*) ≤ 1, and hence

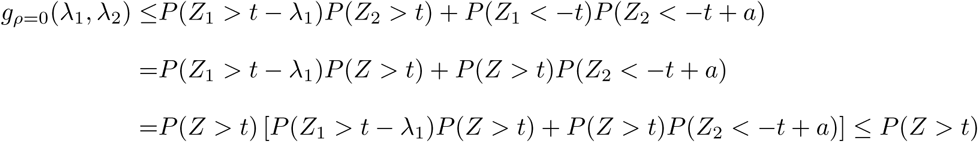

*Case B* When 0 ≤ *t* ≤ *a* ≤ *λ*_1_, we will show *g*_*ρ*=0_(*λ*_1_, *λ*_2_) − [1 − Φ(*t*)] = *g*_*ρ*=0_(*λ*_1_, *λ*_2_) − *P* (*Z > t*) ≤ 0. We first note that

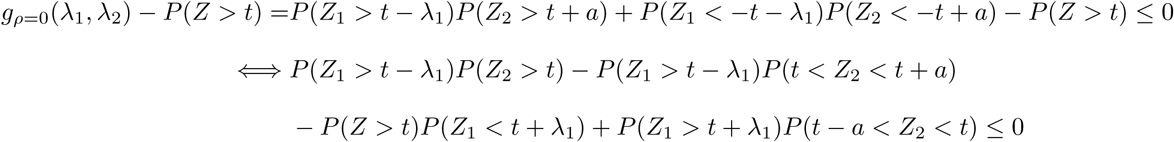

We write the left hand side of the last inequality as *I* + *II*, where

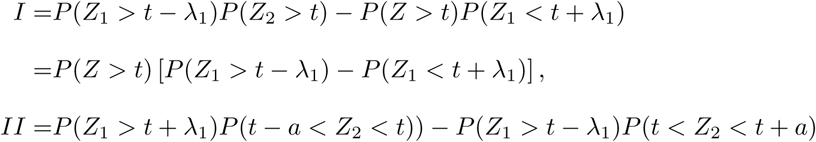

To show *I* + *II* ≤ 0, it suffices to show *I* ≤ 0 and *II* ≤ 0. Since *t >* 0, it is obvious that *I* ≤ 0. To show *II* ≤ 0, it is equivalent to show that 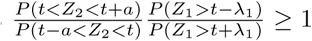. We let 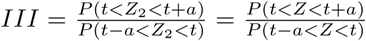 and 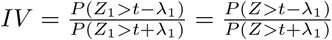. It suffices to show that *III · IV* ≥ 1. *IV* obtains its minimum when *λ*_1_ is minimized. Because 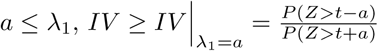. Therefore,

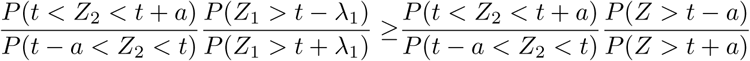

Let 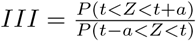 and 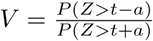. To show that *III · V* ≥ 1, we introduce Lemma 9 and Lemma 10.

#### Lemma 9

(Abramowitz, 1965[12]). 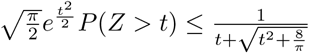 *t* ≥ 0.

#### Lemma 10.

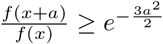 *for t* − *a < x < t < a.*

The proof of Lemma 10 will be provided in Appendix B3.

Let 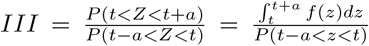. With change of variable *x* = *z − a* in the numerator, 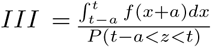. By Lemma 10, 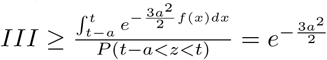.

We know that 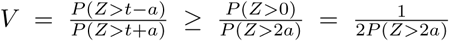. By Lemma 9, 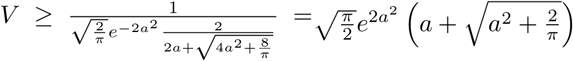.

Therefore, 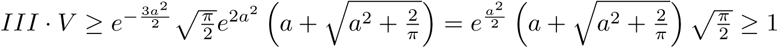, when *a* > 0.

*Case C* When 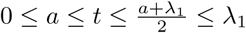, we will show *I* + *II* ≤ 0. To show *I* + *II* ≤ 0, it’s suffice to show that *III · IV* ≥ 1. Recall that 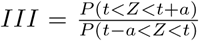 and 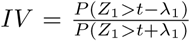. For fixed *t, III* obtains its minimum when *a* is maximized. Given that 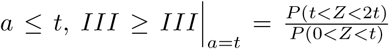. For fixed *a* and *t, IV* obtains its minimum when *λ*_1_ is minimized. Given that the 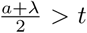, 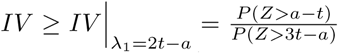. For fixed *t*, this ratio obtains its minimum when *a* is maximized. Given that 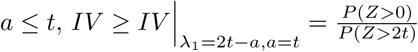.

To show 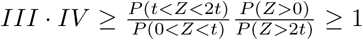, note that 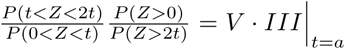 from the *Case B*. Hence, it follows that 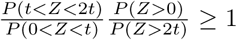.

To summarize, we have shown that for any *t >* 0 and *λ*_1_, *λ*_2_ ∈ Θ_0_, we have *g*(*λ*_1_, *λ*_2_) *< P* (*Z > t*). This completes the proof of Lemma 4 and Theorem 5.

### B2. Proof of Lemma 8

*Proof.* To show *g*(*λ*_1_, *λ*_2_, *ρ*) ≤ *g*(*λ*_1_, *λ*_2_, 0), we will utilize Claims 1 and 2.

*Claim* 1. Suppose *W*_1_ and *W*_2_ are such that

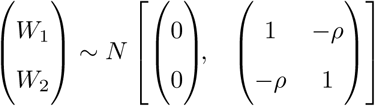

where *ρ >* 0. If *c >* 0, *d >* 0, then *P* (*W*_2_ *> d*|*W*_1_ *> c*) ≤ *P* (*W*_2_ *> d*).

*Proof of Claim 1.* Given the joint distribution of *W*_1_ and *W*_2_ given above, we konw *W*_2_|*W*_1_ ∼ *N* (−*ρW*_1_, 1 − *ρ*^2^). Thus, for *d >* 0, *P* (*W*_2_ *> d*) ≥ *P* (*W*_2_ *> d*|*W*_1_ = *w*_1_) if *w*_1_ *>* 0. It follows that

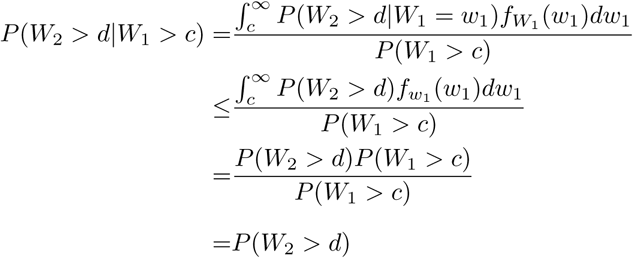

*Claim* 2. For *W*_1_ and *W*_2_ defined above, if *c >* 0 and *d >* 0, then *P* (*W*_2_ *> d*|*W*_1_ *<* −*c*) ≥ *P* (*W*_2_ *> d*). *Proof of Claim 2.*

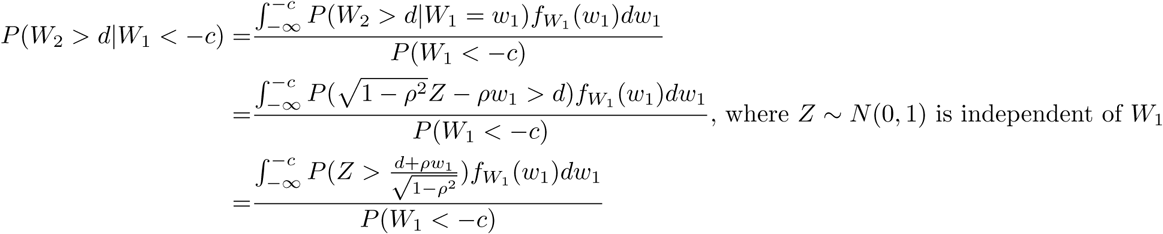

We first show the result for *c* ≥ *d*. In this case, 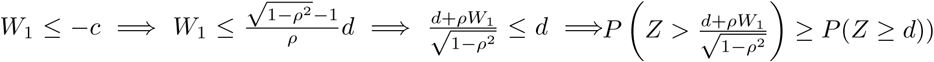. So

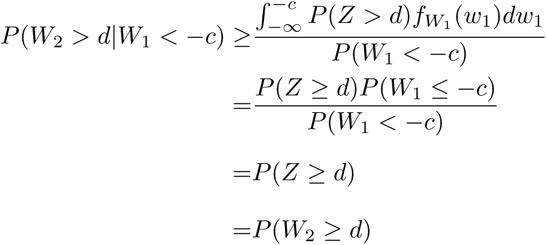

For *c < d*, it suffices to show,

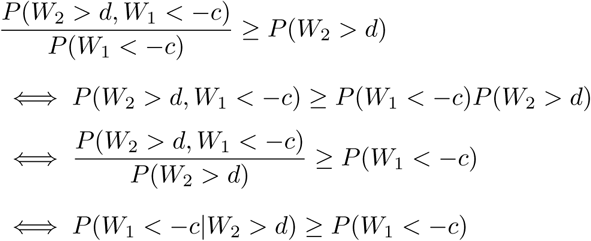

This is equivalent to the result with *c > d*.

Now we go back to the proof of Lemma 8. We discuss three cases based on the magnitude of *t* relative to *a* and *λ*_1_.

*Case 1 Assume* 0 ≤ *a* ≤ *λ*_1_ ≤ *t.*

By Claim 1, we can show that *P* (*Z*_1_ *> t* − *λ*_1_, *Z*_2_ *> t* + *a*) ≤ *P* (*Z*_1_ *> t* − *λ*_1_)*P* (*Z*_2_ *> t* + *a*), and *P* (*Z*_1_ *< t*−*λ*_1_, *Z*_2_ *<* −*t*+*a*) ≤ *P* (*Z*_1_ *> t*+*λ*_1_)*P* (*Z*_2_ *> t*−*a*). Thus, *g*(*λ*_1_, *λ*_2_, *ρ*) = *P*_*ρ*_(*Z*_1_ *> t*−*λ*_1_, *Z*_2_ *> t*+*a*)+*P*_*ρ*_(*Z*_1_ *< t* − *λ*_1_, *Z*_2_ *<* −*t* + *a*) ≤ *P* (*Z*_1_ *> t* − *λ*_1_)*P* (*Z*_2_ *> t* + *a*) + *P* (*Z*_1_ *> t* + *λ*_1_)*P* (*Z*_2_ *> t* − *a*) ≤ *g*(*λ*_1_, *λ*_2_, 0).

*Case 2 Assume* 0 ≤ *a* ≤ *t < λ*_1_.

Since *P* (*Z*_1_ *> t* − *λ*_1_, *Z*_2_ *> t* + *a*) = *P* (*Z*_2_ *> t* + *a*) − *P* (*Z*_2_ *> t* + *a, Z*_1_ *< t* − *λ*_1_). By Claim 2, *P* (*Z*_1_ *> t* − *λ*_1_, *Z*_2_ *> t* + *a*) ≤ *P* (*Z*_2_ *> t* + *a*) − *P* (*Z*_1_ *> t* + *a*)*P* (*Z*_1_ *< t* − *λ*_1_) = *P* (*Z*_2_ *> t* + *a*)*P* (*Z*_1_ *> t* − *λ*_1_). Thus, *P* (*Z*_1_ *> t* − *λ*_1_, *Z*_2_ *> t* + *a*) ≤ *P* (*Z*_2_ *> t* + *a*) − *P* (*Z*_2_ *> t* + *a*)*P* (*Z*_1_ *> t* − *λ*_1_)*P* (*Z*_1_ *< t* − *λ*_1_) = *P* (*Z*_2_ *> t* + *a*).

By Claim 1, *P* (*Z*_1_ *> t* + *λ*_1_, *Z*_2_ *> t* − *a*) ≤ *P* (*Z*_1_ *> t* + *λ*_1_)*P* (*Z*_2_ *> t* − *a*). By an argument similar to Case 1, *g*(*λ*_1_, *λ*_2_, *ρ*) ≤ *g*(*λ*_1_, *λ*_2_, 0).

*Case 3 Assume* 0 ≤ *t < a* ≤ *λ*_1_.

Similar to Case 2, it can be shown that *P* (*Z*_1_ *> t* − *λ*_1_, *Z*_2_ *> t* + *a*) ≤ *P* (*Z*_2_ *> t* + *a*)*P* (*Z*_1_ *> t* − *λ*_1_). Using Claim 2, *P* (*Z*_1_ *> t* + *λ*_1_, *Z*_2_ *> t* − *a*) = *P* (*Z*_1_ *<* −*t* − *λ*_1_) − *P* (*Z*_1_ *<* −*t* − *λ*_1_, *Z*_2_ *>* −*t* + *a*) ≤ *P* (*Z*_1_ *<* −*t* − *λ*_1_) − *P* (*Z*_2_ *>* −*t* + *a*)*P* (*Z*_1_ *<* −*t* − *λ*_1_) = *P* (*Z*_1_ *<* −*t* − *λ*_1_)*P* (*Z*_1_ *> t* − *a*).

Overall, for any *t >* 0, we have shown that *g*(*λ*_1_, *λ*_2_, *ρ*) ≤ *g*(*λ*_1_, *λ*_2_, 0) holds for all *t >* 0. This completes the proof of Lemma 8. By an argument similar to Case 1, *g*(*λ*_1_, *λ*_2_, *ρ*) ≤ *g*(*λ*_1_, *λ*_2_, 0).

### B3. Proof of Lemma 10

*Proof.* 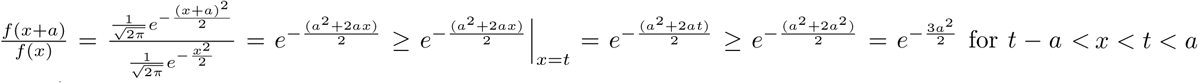.

## Appendix C

In this appendix, we describe two naive procedures in addition to the naive procedure based on 2-sided TSTTs described in Section 2.

### C1. Naive procedure based on 1-sided TSTTs

To test for a strictly monotonic trend against the broad null, the two possible directions of the trend can be tested separately: the strictly increasing trend and the strictly decreasing trend. Each direction can be tested via a 1-sided TSTT between groups 1 and 2, and another such test between groups 2 and 3. For the increasing trend for example, we first test whether *µ*_1_ is significantly smaller than *µ*_2_, and likewise for *µ*_2_ and *µ*_3_. For the decreasing trend, a similar process can be followed. After four 1-sided TSTTs are conducted, we reject the null hypothesis in Equation (1) if for either direction both 1-sided TSTTs result in rejections of their corresponding nulls. Because we have multiple tests, the significance level for each test needs to be adjusted. Using Bonferroni correction, it can be shown that if we conduct each 1-sided TSTT at level 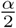, then the overall type 1 error rate is controlled at *α*. Similar with 2-sided TSTTs procedure, the p-value is a linear transform from the greedy test. The p-value for 1-sided TSTTs procedure is *P*_1−*sidedT ST T*_ = max(*p*_1_, *p*_2_). This procedure is also laid out in Table 2.

### C2. Naive procedure based on confidence intervals

The test can be conducted using simultaneous 2-sided confidence intervals for the group means. The confidence interval for each of the sample means can be obtained based on the t-distribution with degrees of freedom equal to the associated sample size minus 1. The broad null hypothesis is then rejected if and only if none of the confidence intervals overlap and the sample means have a monotonic trend. To yield an overall confidence level of 1 − *α*, the Bonferroni method can be used to form the confidence interval for each mean at level 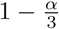. It can be shown that this testing procedure guarantees an overall type 1 error control at *α*.

## Notes

### Competing Interest Statement

The authors have declared no competing interest.

